# Biophysics of object segmentation in a collision-detecting neuron

**DOI:** 10.1101/216333

**Authors:** Richard B. Dewell, Fabrizio Gabbiani

## Abstract

Collision avoidance is critical for survival, including in humans, and many species possess visual neurons exquisitely sensitive to objects approaching on a collision course. The most studied such collision-detecting neuron within the optic lobe of grasshoppers has long served as a model for understanding collision avoidance behaviors and their underlying neural computations. Here, we demonstrate that this neuron detects the spatial coherence of a simulated impending object, thereby carrying out a computation akin to object segmentation critical for proper escape behavior. At the cellular level, object segmentation relies on a precise selection of the spatiotemporal pattern of synaptic inputs by dendritic membrane potential-activated channels. One channel type linked to dendritic computations in many neural systems, the hyperpolarization-activated cation channel, HCN, plays a central role in this computation as its pharmacological block abolishes the neuron's spatial selectivity and impairs the generation of visually guided escape behaviors, making it directly relevant to survival. Our results elucidate how active dendritic channels produce neuronal and behavioral object specificity by discriminating between complex spatiotemporal synaptic activation patterns.

---

Neurons within the brain receive information about the outside world through a continuous ever-changing stream of synaptic inputs. These inputs can arrive thousands of times a second spread out across tens or even hundreds of thousands of different synaptic locations. Ultimately, the primary task of a neuron is to filter out the irrelevant elements of this dynamic stream and extract from the noisy cascade features meaningful for the animal. While the importance of synaptic input timing in this process is well known, due to technical limitations the role of the spatial pattern of dendritic inputs has received less attention. In fact, it is still an unsettled question whether neurons extract information embedded within the broader spatial patterns of ongoing synaptic inputs^1^.

In support of this hypothesis, recent investigations have begun to demonstrate spatial patterning of excitatory and inhibitory synaptic inputs^2,3,4^ and dendritic processes capable of discriminating between different such patterns^5,6,7^. For instance, local synaptic clustering can produce supralinear summation which enhances the selectivity of visual neurons^5,7^. Dendritic spikes and NMDA receptors can amplify local patterns of synaptic inputs, conferring directional selectivity to some retinal ganglion cells^8,9^. While recent results have begun to illustrate the functional role of fine scale synaptic patterning in many neurons^10,11,12,13^, whether they also discriminate broader spatiotemporal patterns consisting of thousands of synaptic inputs remains unaddressed. The nonlinear dynamics employed within dendrites for this purpose will likely depend on the spatiotemporal statistics of presynaptic activities and the neuronal function that needs to be computed^14,15^. To address these issues, we focus on large scale processing of synaptic inputs and the dendritic computations required for visual object segmentation in the context of collision avoidance behaviors.

The spatiotemporal sequence of synaptic inputs relevant to collision avoidance is determined by the statistics of the approaching object. Objects approaching on a collision course or their simulation on a screen, called looming stimuli, produce a characteristic visual stimulus on the observer's retina, expanding coherently in all directions with increasing angular velocity. Discriminating this retinal pattern from that of optic flow or from that of an object approaching on a miss trajectory requires integrating information across many points in time and space. Among neurons capable of such discrimination^16,17,18,19,20^, the best understood is the lobula giant movement detector (LGMD, **Fig. 1a**), an identified neuron of the grasshopper optic lobe located three synapses away from photoreceptors^21^. The LGMD responds maximally to looming stimuli^22,23,24^ with a characteristic firing rate profile^24,25^ (**Fig. 1b**) that has been tightly linked to initiating escape behaviors^26^. This characteristic firing profile is maintained even when an approaching stimulus is embedded in a random motion background, suggesting that the LGMD may be able to effectively segment visual objects^27,28^. This sensory selectivity and behavioral relevance has led to the LGMD serving as a model for neural computations involved in the visual detection and avoidance of impending collision^29^.

**Figure 1.**
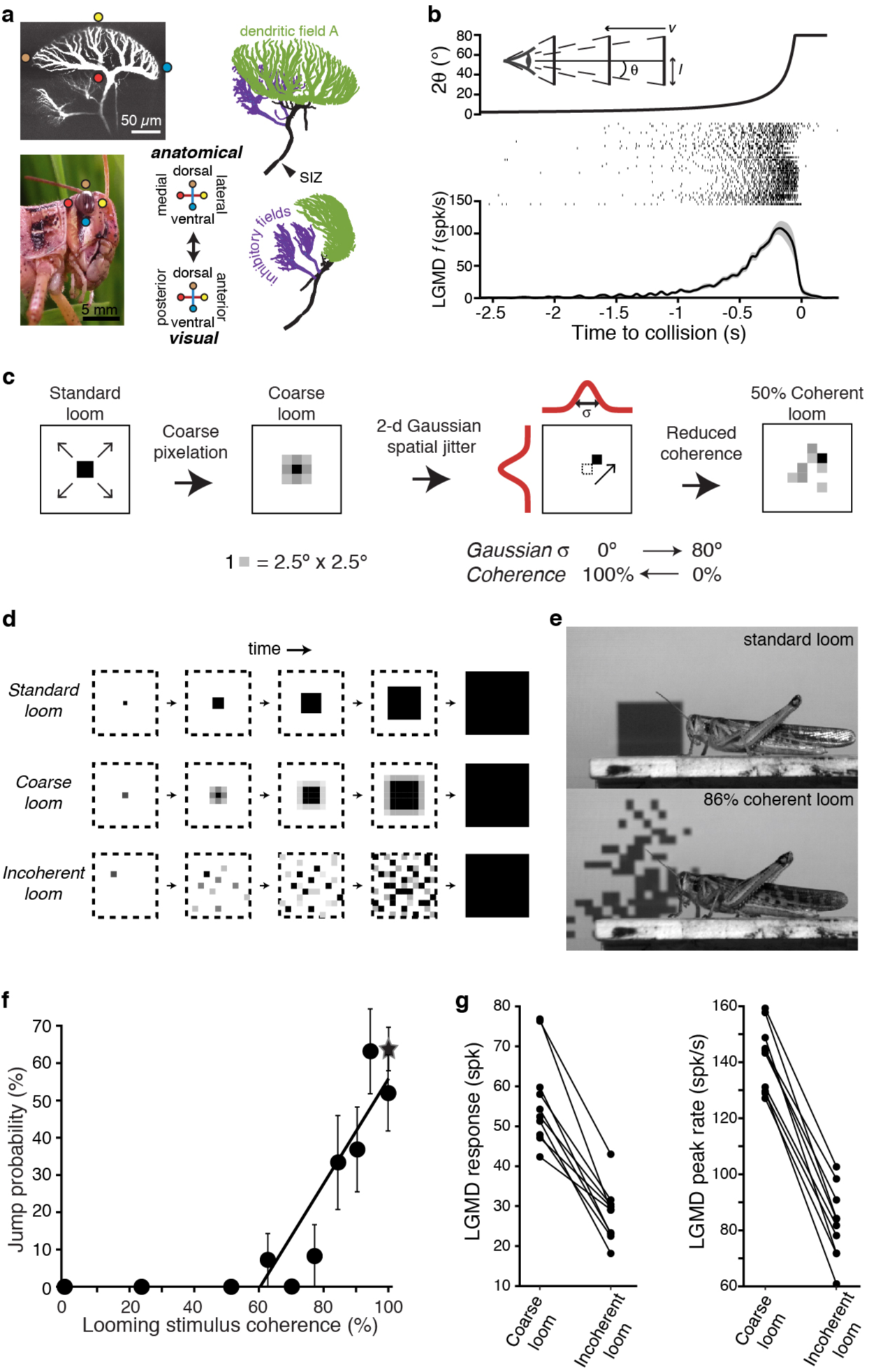
LGMD responses and escape behavior are sharply tuned to the spatial coherence of looming stimuli. (**a**) LGMD 2-photon scan (top, left), eye close-up of *Schistocerca americana* (bottom, left), rostral, and lateral view of a LGMD reconstruction used for modeling (top and bottom right). Excitatory dendritic field in green, SIZ: spike initiation zone. Colored dots illustrate the retinotopic mapping of excitatory inputs to the LGMD. (**b**) Top, schematic of visual stimulus, half-size *l*, approach speed *v*, half-angular subtense at the eye, *θ*. Note the non-linear increase in angular subtense (2*θ*), characteristic of looming stimuli. Middle, spike rasters of the LGMD responses to looming stimuli. Bottom, mean instantaneous firing rate (*f*) of LGMD looming response. Shaded area is ±1 sem. (c) The coherence of looming stimuli was altered by first applying a coarse pixelation to create photoreceptor sized pixels. Then, a zero-mean random shift was added to the position of these coarse pixels to generate the reduced coherence stimuli. The variance of the random shifting (in degrees) determined the reduction in coherence. (**d**) Illustration of coherent and incoherent stimuli. For coarse looms (middle row) grayscale levels are set so that luminance in each coarse pixel is equal to that of standard looms in every frame. For reduced coherence looms (bottom), the spatial loCa_T_ions of the coarse pixels were altered. (**e**) Video frames from presentation of standard looming (top) and 86% coherent (bottom) stimuli. (**f**) Jump probability increased sharply with stimulus coherence above 50% (r = 0.91, p = 5.9·10^−4^), 202 trials from 66 animals. (**g**) The LGMD's spike count (p = 2.5·10^−4^, Wilcoxon rank sum, WRS) and peak firing rate (p = 1.9·10^−4^, WRS) were lower for 0% coherent than 100% coherent looming stimuli (N=10).

Synaptic inputs onto the LGMD are physically segregated into three dendritic fields, two of which receive non-retinotopically organized inhibitory inputs^30^. The third one, dendritic field A, receives excitatory inputs originating from each ommatidium (facet) on the ipsilateral compound eye in a precise retinotopic projection^31,32,33^ (**Fig. 1a**). These excitatory synaptic inputs to the LGMD are segregated by ommatidia and arranged in columnar fashion over an entire visual hemifield, so the LGMD's dendritic arbor has unique access to the entire spatial visual pattern activated by an approaching stimulus. Like in cortical neurons that often receive inputs from tens of thousands of synapses spread across their dendritic arbors, little is known on whether the LGMD detects the spatial patterning of its synaptic input. The precise retinotopy of field A^32,33^ offers the possibility to experimentally control this patterning in vivo over a broad, natural range by changing the spatial aspect of visual stimuli, and thus test the LGMD’s ability to discriminate spatial patterns consisting of thousands of synaptic inputs. Here, we demonstrate that the LGMD indeed discriminates the spatial coherence of approaching objects, that hyperpolarization-activated cyclic nucleotide-gated nonselective cation (HCN) channels within the retinotopic dendrites are critical to this discrimination through complex interactions with other membrane channels, and that this neural selectivity enhances the animal's ability to effectively avoid approaching predators.

## RESULTS

### Tuning of the LGMD and escape behavior to stimulus coherence

To control the stimulus pattern, as in our earlier work^34^, we replaced standard looming stimuli by equivalent ones pixelated at the spatial resolution of photoreceptors on the retina, called ‘coarse’ looming stimuli (**Fig. 1c,d**). The LGMD responds equally to standard and coarse looming stimuli^34^. We could then alter the coherence of these stimuli with minimal change to the temporal pattern of activation experienced by individual photoreceptors by adding a random spatial jitter to each ‘coarse pixel’ (**Fig. 1c**). Spatial stimulus coherence was varied from random (0%) to perfectly coherent (100% = standard or coarse looming stimulus; **Materials and Methods**; **Supplementary Videos 1 and 2**). Looming stimuli with full and reduced coherence were presented to unrestrained animals while recording the probability of escape jumps (**Fig. 1e and Supplementary Video 3**). Locusts showed a strong behavioral selectivity to spatial coherence; stimuli with less than 50% spatial coherence elicited no escape jumps, but jump probability increased rapidly with coherence above 50% (**Fig. 1f**). The firing rate of the LGMD was also highly sensitive to stimulus coherence, with sharply reduced spike count and peak spike rate at 0% coherence (**Fig. 1g**). This demonstrates that the spatial coherence of an approaching object determines both the LGMD's response and the animal's decision of whether to escape.

### HCN channels in dendritic field A are responsible for coherence tuning

None of the known properties of the LGMD or its presynaptic circuitry could explain this spatial selectivity. Previous experiments showed that the strength of excitatory inputs encodes the temporal characteristics of the approaching object independent of its spatial pattern^34^. Additionally, simulations confirmed that spatial clustering of synaptic inputs that occurs with coherent stimuli would reduce summation in passive LGMD dendrites^32^, as further elaborated below. The LGMD's selectivity for the spatial characteristics of an approaching object must therefore be determined by active processing within the dendrites of field A. No active conductances, however, have yet been characterized within these dendrites. Evidence suggests that neither the fast Na^+^ nor Ca^2+^ channels that produce supralinear summation in many other neurons are present there^34,35^. Since in many neural systems HCN channels have been found to influence dendritic computations, and previous experiments suggested putative HCN channels within the LGMD^36^, we hypothesized that HCN channels within field A might be involved in discriminating the spatial coherence of approaching objects.

To test for the presence of HCN channels we used current and voltage steps, as well as application of known channel blockers and modulators^37^, during visually guided recordings from each of LGMD's three dendritic fields and near the spike initiation zone (SIZ; **Fig. 2a**). Hyperpolarization produced a characteristic rectifying sag, that was abolished by the HCN channel blockers ZD7288 and Cs^+^ (**Supplementary Fig. 1a-c**). At the same peak hyperpolarization (**Fig. 2b**, Methods), the HCN channels’ conductance (gH) produced a larger, faster sag in field A (**Fig. 2b-d**), consistent with HCN channels being localized there and the current passively propagating to the rest of the LGMD. Recordings at different locations within dendritic field A also revealed an increase in sag with distance from the SIZ (**Fig. 2e**) suggesting a higher channel density distally in field A. To characterize the channel kinetics, field A dendrites were voltage clamped revealing an activation curve (**Fig. 2f**) and time constant (**Fig. 2g**) similar to that of HCN2 channels^37^. Application of cAMP shifted the activation curve of g_H_ (**Fig. 2f**), resulting in increased activation at rest, while ZD7288 application abolished g_H_ activation (**Fig. 2h**) consistent with results in other systems.

**Figure 2.**
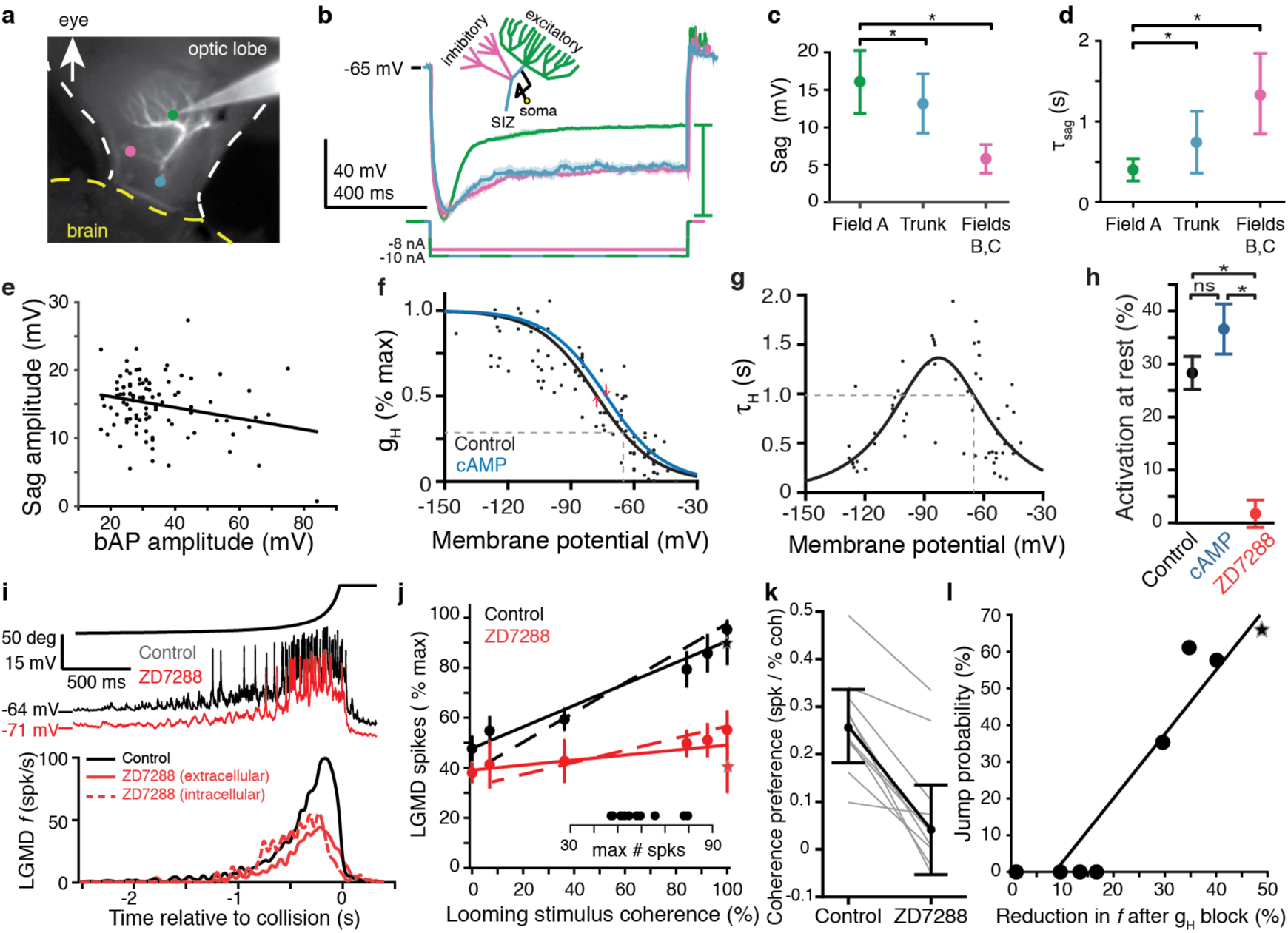
HCN channels in dendritic field A responsible for spatial coherence sensitivity. (**a**) Image showing the LGMD stained *in vivo* with Alexa 594 and the recording electrode with tip at the green dot. Colored dots indiCa_T_e the recording loCa_T_ions of traces shown in (**b**). (**b**) Schematic of the LGMD’s dendritic subfields and example traces showing larger rectifying sag in field A than in field C or near the SIZ. Solid lines are the average response with shaded region of ± 1 sd. Sag amplitude was measured as the amount of rectifiCa_T_ion from peak hyperpolarization to steady state, as indiCa_T_ed by the green bar. Responses with nearly identical peak hyperpolarization were obtained by adjusting the injected current steps (-10 nA in field A and near the SIZ; -8 nA in field C). (**c**) Sag amplitudes following steps from rest yielding peak hyperpolarizations between −95 and −115 mV are consistently larger in field A (N=82,58) than in the trunk (N=11,13) or inhibitory subfields (N=6,6; p < 0.001, KW-MC). (**d**) The sag time constant for these responses was smaller in field A (N=82,58) than in the trunk (N=11,13) or inhibitory subfields (N=6,6; p < 0.01, KW-MC). For **c,d**, bar heights are median and error bars are 1 mad. (**e**) Sag amplitude decreased with increased backpropagating action potential (bAP) amplitude, a measure of distance from the spike initiation zone (r = -0.25, p = 0.01, N=104,69). (**f**) Activation curve of g_H_ measured in voltage clamp. Black line is control (N=8,7; *v*_*1/2*_ = −78 mV, 28% of max at RMP, dashed line; R^2^ = 0.69) and blue line after local appliCa_T_ion of cAMP (N=6,6; *v*_*1/2*_ = −73 mV, 35% of max at RMP). Red arrows indiCa_T_e shift in *v*_*1/2*_. (**g**) Time constant of g_H_ from voltage clamp recordings (N=8,7; τ_H_ max = 1.34 s, at −83 mV; τ_H_ = 985 ms at RMP, dashed line; steepness = 20 mV; R^2^ = 0.61). (**h**) Resting activation of HCN channels, relative to max, displayed as median and mad (control N=82,58; cAMP N=6,6; ZD7288 N=13,10; *: p < 0.001, ns: p = 0.076, KW-MC). (**i**) Intracellular recordings of LGMD’s membrane potential in response to looming stimuli show decreased RMP and activation after blockade of g_H_ (top). Bottom, mean instantaneous firing rates (*f*) in response to looming stimuli declined after intra- or extra-cellular appliCa_T_ion of ZD7288 (N=10,10 p = 4.1·10^−5^, WRS). (**j**) Responses increased with stimulus coherence in control conditions (r = 0.83, p = 2.9·10^−14^), but after g_H_ block the coherence-dependent increase was removed (p = 2.1·10^−6^, ANCOVA test of slopes, N=10,10). Solid lines and dots are coarse loom data, stars are standard loom, error bars are ± 1 sd, and dashed lines are compartmental simulation results. Insets show plot normalization values. (**k**) For all experiments coherence preference decreased after g_H_ blockade (N = 10, gray lines). Coherence preference was calculated as the increase in spike count per percent increase in stimulus coherence. Black dots and lines show the average coherence preference decreased by 0.19 spikes per percent stimulus coherence (p = 1.3·10^−4^, paired t-test). (**l**) Jump probability for a stimulus correlates strongly with its g_H_-dependent increase in firing (r = 0.94, p = 4.1·10^−4^). **N: number of recordings, number of animals.**

To examine whether these HCN channels could be responsible for spatial discrimination, we presented visual stimuli before and after their pharmacological blockade. Responses to looming stimuli were reduced by 61% after HCN blockade within field A (**Fig. 2i**). Visual responses to localized luminance transients, however, were similar before and after blockade of g_H_ (**Supplementary Fig. 2**). After ZD7288 blockade, responses at all coherence levels were like those at zero coherence before block (**Fig. 2j**; un-normalized, individual data in **Supplementary Fig. 3a**). As explained below, this change in selectivity was reproduced by a biophysical model of the LGMD (**Fig. 2j**, dashed lines). For each experiment, we defined coherence preference as the slope of the linear fit to the number of spikes fired by the LGMD as a function of stimulus coherence (**Supplementary Fig. 3a**). For every animal tested, the coherence preference was reduced after g_H_ blockade, decreasing from a median of 0.26 to 0.04 spikes per percent coherence (**Fig. 2k**). To ensure that these effects were intrinsic to the LGMD, we ascertained that blocking g_H_ with Cs^+^ also reduced the coherence selectivity (**Supplementary Fig. 3b**; **Materials and Methods**). Comparing the jump probabilities at each coherence level with the gH-dependent increase in firing revealed a strong correlation (**Fig. 2l**). Furthermore, responses to faster looming stimuli, which fail to produce escape behaviors before the projected time of collision^38^, showed a smaller gH-dependent increase in firing (**Supplementary Fig. 3c**). Therefore, g_H_ increased responses specifically to stimuli which evoke escape, suggesting that the gH-dependent enhancement produced the increase in escape selectivity.

**Figure 3.**
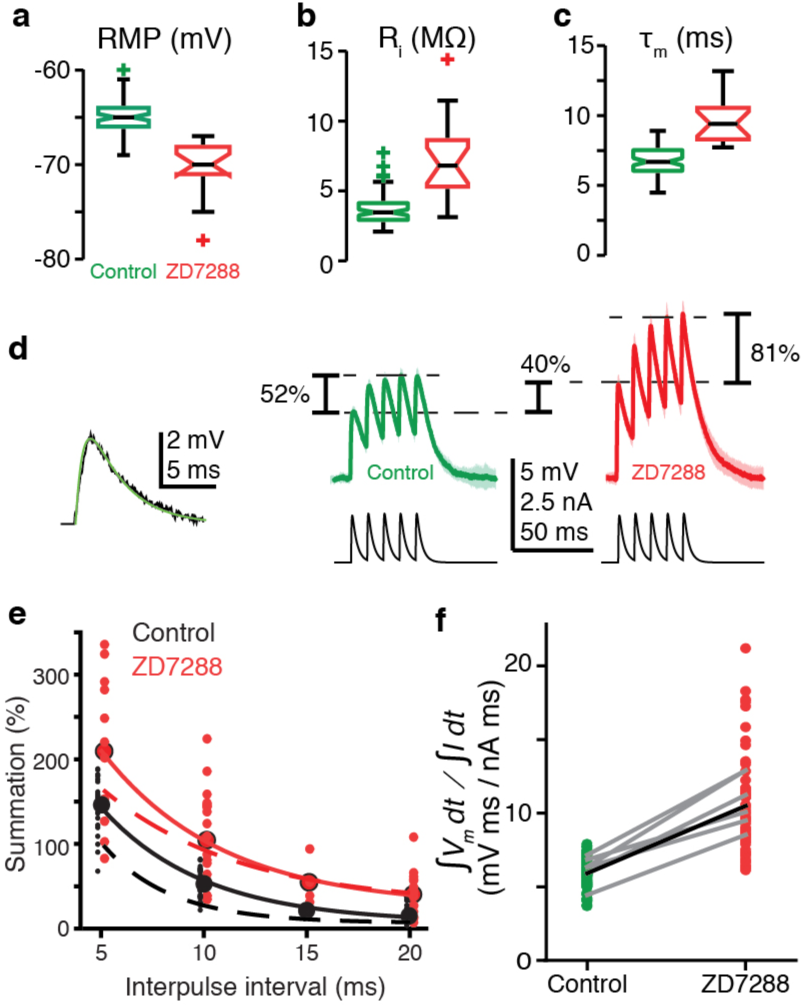
g_H_ conductance decreases EPSP amplitudes and summation. (**a**) RMP in field A decreased after blockade of g_H_ by ZD7288 (control N=82,58; ZD7288 N=15,9; p = 1.9·10^−9^, WRS). (**b**) Input resistance (ri) in field A increased after g_H_ blockade (control N=78,58; ZD7288 N=17,10; p = 6.1·10^−7^, WRS). (**c**) Membrane time constant (τ_m_) also increased in field A after g_H_ block (control N=82,58; ZD7288 N=16,10; p = 1.3·10^−8^, WRS). (**d**) Left, example visual EPSP (black) and sEPSP (green). right, example responses to a series of 5 sEPSPs with 10 ms interpulse interval. On average ZD7288 block of g_H_ led to 51% increase in 1^st^ sEPSP amplitude (p = 6.0·10^−4^, WRS) and subsequent summation of sEPSPs from 53% in control to 104% (p = 0.001, WRS) control N=14,7; ZD7288 N=9,5. (**e**) Summation increased after g_H_ removal for all interpulse intervals. Points and solid lines show experimental data, dashed lines are simulation results. (**f**) The integrated membrane potential (V_m_) normalized by the integrated current (total charge) increased an average of 77% after ZD7288 (thick black line; p = 1.7·10^−19^, WRS, N =14,7 in control and 9,5 in ZD7288). Gray lines are from 6 recordings held through puffing (p <= 0.01, paired t-test).

### HCN channels affect membrane properties and synaptic summation

How HCN channels could impart this selective enhancement is not obvious. In no neurons have HCN channels been shown to increase summation of spatially coherent inputs, and often g_H_ has net inhibitory effects^37,39^. To determine how HCN channels produced the selective enhancement of looming responses, we investigated the effects of g_H_ on membrane excitability within field A. g_H_ increased the resting membrane potential (RMP) by ~6 mV in field A, which would bring the neuron closer to spike threshold (**Fig. 3a**). Blockade also revealed g_H_ to decrease input resistance by 50% and the membrane time constant (τ_m_) by 30% (**Fig. 3b,c**), which should substantially reduce the summation of excitatory postsynaptic potentials (EPSPs), as occurs after g_H_ blockade in cortical pyramidal neurons^40,41^. Injecting simulated EPSPs (sEPSPs; **Fig. 3d**) confirmed this reduced summation. After g_H_ blockade, summation from the first to fifth sEPSP increased for all tested delays (**Fig. 3e**; the dashed lines are from a biophysical model, see below), and the integrated sEPSP, normalized by the integrated current increased by 77% (**Fig. 3f**). Neither before nor after g_H_ blockade was supralinear summation ever seen in LGMD dendrites. We verified that the effects of g_H_ were not simply due to a shift in the RMP by hyperpolarizing the LGMD during visual stimuli. Lowering the RMP without altering input resistance and τ_m_ produced less reduction in coherence preference than blocking gH, with responses to standard looming stimuli reduced by 20%. (**Supplementary Fig. 3d**). The mix of excitatory and inhibitory electrotonic effects of g_H_ does not provide any simple explanation for the large enhancement in looming responses or the conveyed coherence selectivity.

### K^+^ channels complement HCN channels in generating coherence tuning

That g_H_ increased looming responses twofold despite decreasing EPSP amplitude and summation by half appears counterintuitive. To explore this apparent contradiction, we investigated interactions between HCN and other dendritic channels. In several systems, HCN channels have indirect excitatory effects through inactivation of co-localized voltage-gated K^+^ channels^41,42,43,44^. To test whether this was also the case in dendritic field A of the LGMD, we measured visual responses in the presence of 4-aminopyridine (4AP), a blocker of inactivating K^+^ channels^45^. Application of 4AP, either intracellularly or extracellularly, increased the resting membrane potential in field A by 2-5 mV and increased responses to all visual stimuli, but responses to looming stimuli increased the least (**Fig. 4a,b**; un-normalized, individual data in **Supplementary Fig. 3e**). The complementary effects of HCN and K^+^ channels was best revealed by plotting the relative changes in looming responses after their block by ZD7288 and 4AP, respectively (**Fig. 4c**). Thus, while HCN channels predominantly boosted responses to coherent stimuli, K^+^ channels mainly decreased responses to incoherent ones. This increase in incoherent responses after blocking K^+^ channels was also reproduced in biophysical simulations (**Fig. 4b,c** dashed lines; see below).

**Figure 4.**
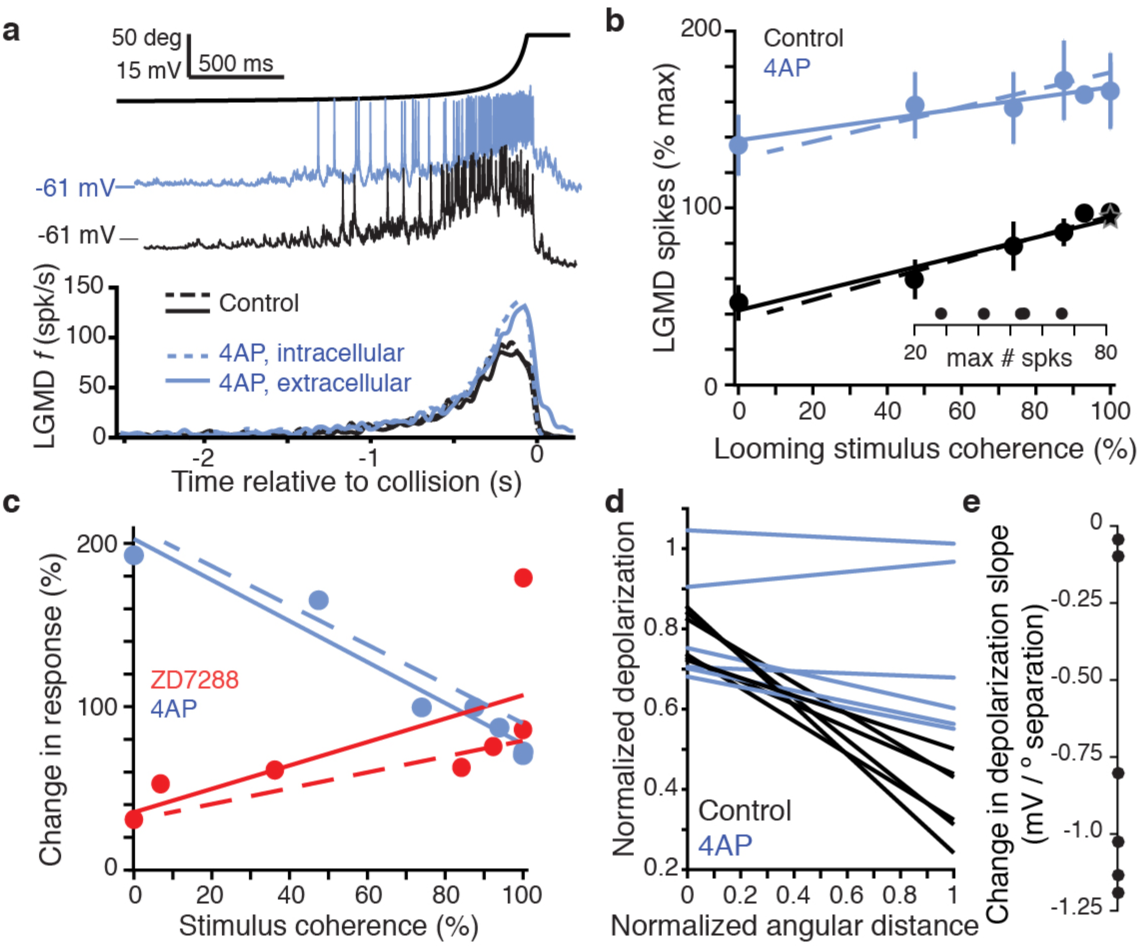
A 4AP-sensitive K^+^ conductance decreases responses to incoherent stimuli. (**a**) The time course of the looming stimulus is indiCa_T_ed on top by its subtended visual angle (2*θ*, see **Fig. 1b**) and (middle) example recordings of LGMD membrane potential in response to looming stimuli before and after local puff of 4AP. Below, average firing rate of the LGMD before and after appliCa_T_ion of 4AP. Both intracellular and extracellular appliCa_T_ion had the same effect on looming responses (N=5,5 for extracellular, N=3,3 for intracellular). (**b**) 4AP increased responses to all stimuli, and reduced the coherence-dependent increase in firing (r = 0.83 p = 8.4·10^−9^ for control, r = 0.36 p = 0.05 after 4AP, N=5,5). Plotted as in **Fig. 2j**. (**c**) Comparison of the relative effects of ZD7288 (r = 0.395, p = 4.1·10^−3^) and 4AP (r = -0.58, p = 6.3·10^−4^). Dashed lines in **b** and **c** are simulation data. (**d**) Membrane potential changes showed a strong negative correlation with distance between stimulated regions in control (r = -0.54, p = 4.1·10^−8^), but 4AP appliCa_T_ion significantly reduced this effect (p = 0.006, ANCOVA test of slopes, N=7,7) Each line pair is from a different time interval of the stimulus. (**e**) Within each time window, the 4AP sensitive current caused a decrease in response to spatially distant inputs (p = 0.019, paired t-test, N=7,7).

To further confirm that these K^+^ channels were exerting a spatially-dependent effect on synaptic integration, we recorded intracellularly from dendritic field A before and after 4AP application while presenting looming stimuli with varying degree of coherence. We measured the dendritic membrane potential over multiple time windows during the stimuli and compared it with the distance between currently changing coarse pixels and those pixels already fully darkened (**Supplementary Fig. 4a-f**). Spatially dispersed excitatory inputs produced less depolarization of the dendrites in control conditions, but this relationship was greatly reduced after application of 4AP, as summarized in **Fig. 4d**. We quantified the change in depolarization caused by currently changing coarse pixels as a function of the angular distance to fully darkened ones by computing the slope of the linear fits between these two quantities (**Supplementary Fig. 4g**). For each time window, 4AP increased the depolarization produced in field A by more distant stimuli with an average slope difference of 0.72 mV per degree of visual separation (**Fig. 4e**). These experiments confirm that inactivating K^+^ channels selectively decrease responses to spatially dispersed inputs in dendritic field A.

### Compartmental modeling highlights role of K^+^ and Ca^2+^ channel inactivation in coherence tuning

Detailed biophysical modeling was employed to further understand the biophysical mechanisms by which HCN and inactivating K^+^ channels allow the LGMD to discriminate spatiotemporal input patterns based on coherence. First, we confirmed that a model of the LGMD with passive dendrites generated a smaller response to retinotopically arranged looming inputs than the same inputs impinging on random dendritic locations (**Fig. 5a**). This illustrates why implementing coherence preference over an extended dendritic arbor is nontrivial: spatially distributed excitatory inputs produce less reduction in driving force, thus generating a larger current from the same synaptic conductance. Adding HCN channels to the dendrites of this model, in agreement with experimental data, also resulted in stronger responses to spatially scrambled inputs (**Fig. 5b**). As suggested by the results of **Fig. 4**, the subsequent addition of inactivating K^+^ channels in dendritic field A reduced responses to the spatially scrambled inputs, bringing the model in broad agreement with experimental findings (**Fig. 5c**).

**Figure 5.**
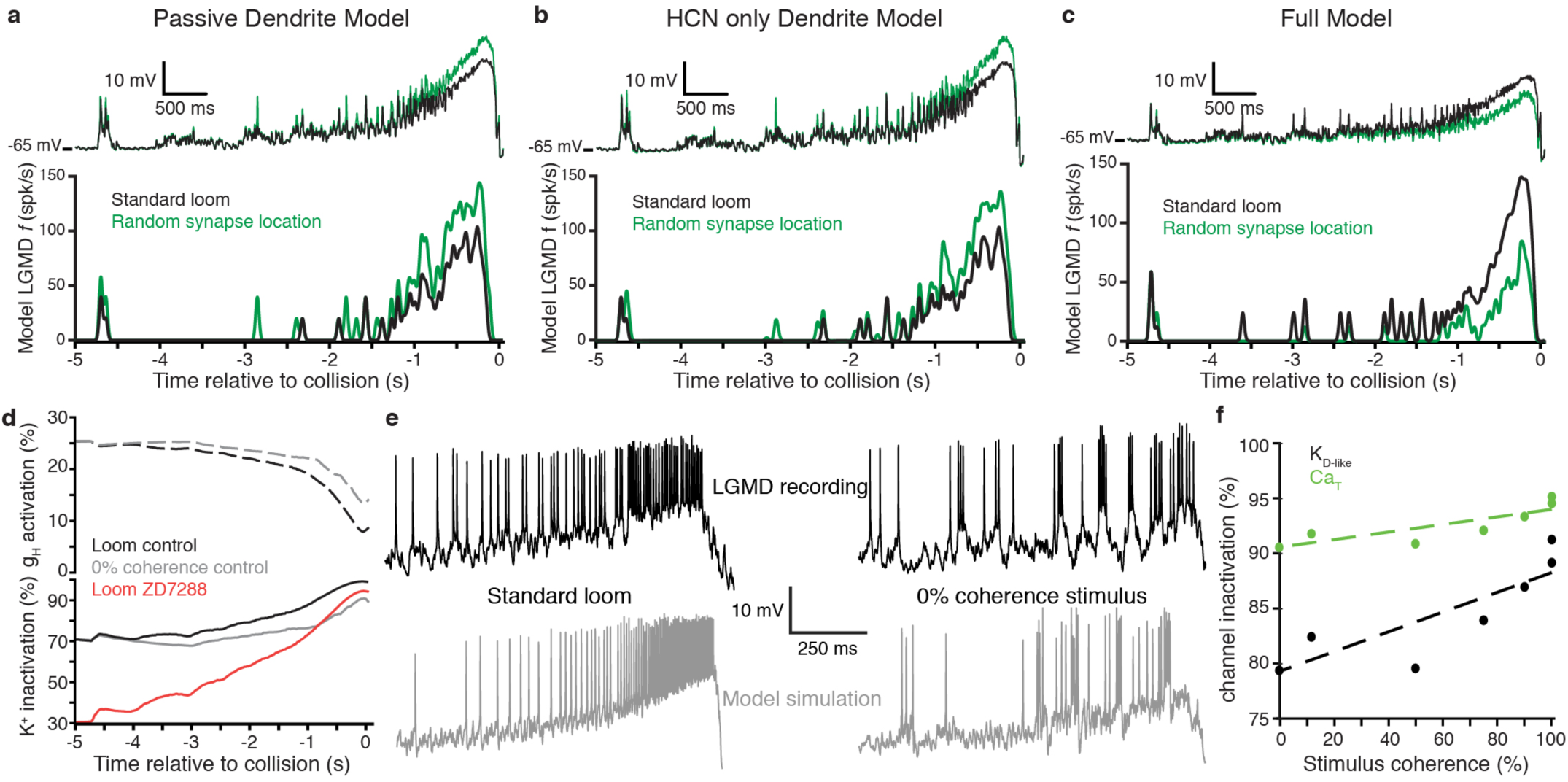
An active biophysical model reproduces the preference for coherent synaptic inputs. (**a**) A model LGMD with realistic morphology and passive dendrites generated a stronger response to spatially randomized inputs (green) than retinotopically arranged ones (black). Spatially randomizing inputs increased mean membrane potential 1.23 ± 0.01 mV at the base of field A (top) and increased firing by 59% (bottom). (**b**) Adding HCN channels to the dendrites did not change this trend. Spatially random inputs (green) increased mean membrane potential 1.23 ± 0.01 mV at the base of field A (top) compared to spatially coherent ones (black) and increased firing by 64% (bottom). (**c**) The full model, with both HCN and inactivating K^+^ channels, however, had 1.25 ± 0.01 mV lower membrane potential at the base of field A and 61% less spiking in response to spatially random inputs. (**d**) At bottom, the time course of K^+^ channel inactivation shows higher inactivation during a coherent looming stimulus (black) than an incoherent coarse stimulus (gray). After blocking HCN channels (red), the resting inactivation is much less and never reaches the inactivation level of control. At top, HCN channel activation is lower during coherent stimuli since K^+^ channel inactivation leads to increased depolarization. (**e**) Comparison of the membrane potential near the base of field A for experimental data (top) and model simulation (bottom) reveal a steady ramp up in firing rate in response to coherent looming stimuli and a much burstier firing pattern in response to a 0% coherence coarse stimulus. (**f**) The mean channel inactivation during the last 2 seconds before collision increased with stimulus coherence for both K^+^ and Ca_T_ channels.

More precisely, this model reproduced many key experimental results, including the LGMD's preference for spatially coherent inputs and the reduction of this preference after block of g_H_ (**Fig. 2j**); the electrotonic and summation effects of g_H_ (**Fig 3e; Supplementary Fig. 5a-c**); the coherence-dependent increase in firing caused by blocking the inactivating K^+^ channels and their role in suppressing responses to incoherent stimuli (**Fig. 4b,c**). In the model, the inactivating K^+^ channel activity was similar to the KD current that has been hypothesized to influence dendritic integration in pyramidal and Purkinje neurons based on its activity at rest, influence on subthreshold integration within field A dendrites, its apparent slow inactivation, and its 4AP sensitivity^45,46^. We thus call it K_D-like_. During looming stimuli, as inputs continue to impinge nearby for a prolonged period, the K^+^ channels inactivate (**Fig. 5d**, solid black line). During spatially incoherent stimuli, however, the channels across the arbor undergo less inactivation (**Fig. 5d**, solid gray line). The reduction in resting membrane potential that occurs after HCN blockade reduces the resting K^+^ channel inactivation (**Fig. 5d**, red line) so that even with spatially coherent inputs the channels never fully inactivate (**Supplementary Fig. 5e** shows the K_D-like_ activation variable for each stimulus condition). The increased inactivation when HCN channels are present for coherent stimuli produced lower overall conductance of the K_D-like_ channels (**Supplementary Fig. 5f**). HCN activation decreased as channels closed because of dendritic depolarization during looming stimuli (**Fig. 5d**, dashed black line). During incoherent stimuli, there was less dendritic depolarization and less closing of HCN channels (**Fig. 5d**, dashed gray line).

In addition to the dendritic channels, the model also included low-threshold Ca^2+^ channels (CaT) and Ca^2+^-dependent K^+^ channels (K_Ca_) near the SIZ that allowed the LGMD model to fire in bursts^36,35^. In both experimental data and simulations, responses to spatially coherent stimuli generated more sustained, non-burst firing than transient burst firing (**Fig. 5e**; **Supplementary Fig. 5d,e**). The model reproduced the trends in the data qualitatively rather than quantitatively (see Discussion). The decrease in bursting for coherent stimuli was also dependent on Ca_T_ channel inactivation. Coherent stimuli produced a steady ramp up of membrane potential increasing Ca_T_ inactivation, while incoherent stimuli produced more sudden depolarization producing bursts. During the last 2 s before collision, when most firing occurred, the average inactivation of both Ca_T_ and K_D-like_ channels increased with stimulus coherence (**Fig. 5f**).

### HCN channels mediate coherence tuning of escape behaviors

Having found that gH-dependent increase in firing is strongly correlated with jump probabilities (**Fig. 2l**), we sought a more direct test of the hypothesis that g_H_ within the LGMD played a critical role in the animals’ escape from approaching objects. So, we blocked g_H_ in the LGMD in freely behaving animals (**Materials and Methods**). As a further control, we also developed a chronic recording technique allowing us to monitor the descending LGMD output during escape behaviors before and after g_H_ blockade.

Blocking g_H_ in the LGMD reduced escape behavior by 53% for standard looming stimuli compared to saline injection (**Fig. 6a**, left two bars). The coherence preference was also removed by blockade of g_H_: standard looming stimuli no longer produced a higher percentage of escape than reduced coherence stimuli (**Fig. 6a**, red bars). This provides confirmation that the spatial selectivity of synaptic input patterns conferred to the LGMD by the active conductances within its dendrites directly influences the selectivity of escape behaviors. That these behavioral changes were caused by g_H_ blockade within the LGMD was further confirmed by examination of the LGMD's firing pattern. g_H_ blockade by ZD7288 decreased responses to both standard looming stimuli and 89% coherent stimuli (**Fig. 6b,c**). The reduction in firing in the freely moving animals was less than that in the restrained preparation (36% and 60%, respectively), which might be due to an incomplete block of g_H_ after stereotactic injection compared to visually guided puffing (see **Materials and Methods**) or differences in arousal state. To test this, we used the stereotactic injection procedure in restrained animals and saw a 56% reduction in looming responses (**Fig. 6d**) suggesting the difference in firing rate change was most likely due to difference in behavioral state. Our ability to produce a direct change in behavior of freely moving animals from targeted blockade of g_H_ within the LGMD was confirmed by simultaneous extracellular recordings revealing a firing rate change resembling that of intracellular drug appliCa_T_ion, verifiCa_T_ion that the surgical procedures did not reduce the response, and postmortem anatomical verifiCa_T_ion that drug appliCa_T_ion occurred within the region encompassing LGMD’s dendrites (**Supplementary Fig. 6**).

**Figure 6.**
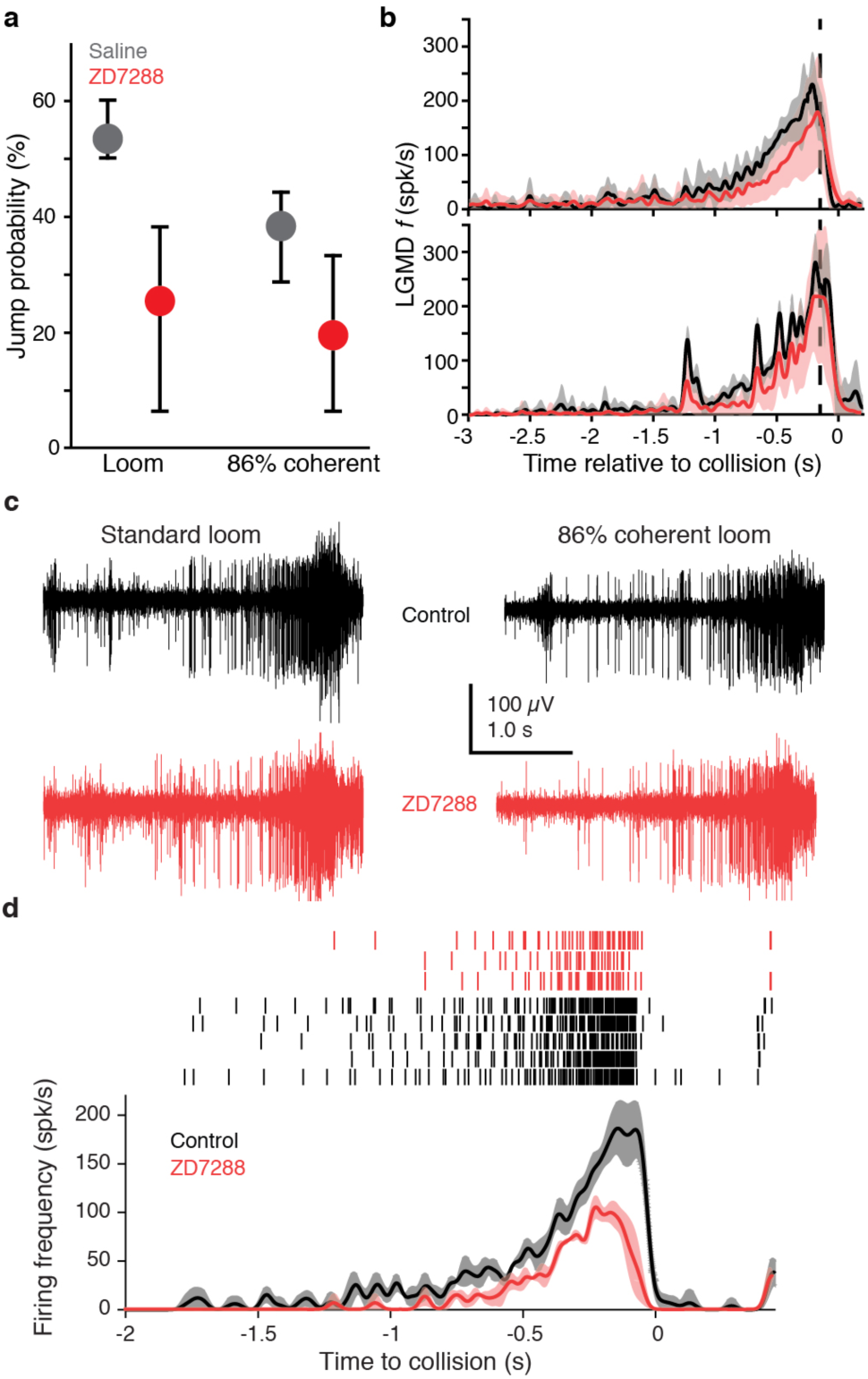
Blocking HCN channels removed coherence preference of escape behavior. (**a**) Jump probability for coherent looming stimuli decreased after injecting ZD7288 into lobula, compared to saline injection (p = 0.008, ASL). Error bars are bootstrapped 95% confidence intervals. Responses to 86% coherent stimuli after ZD7288 injection were not significantly different from responses after saline injection (p = 0.08, ASL) or standard looming responses after ZD7288 injection (p = 0.34, ASL). Saline injection: 48 trials from 5 animals, ZD7288 injection: 41 trials from 5 animals. (**b**) LGMD instantaneous firing rates (*f*) during jump experiments decreased after ZD7288 injection. ZD7288 decreased responses to both stimuli (p = 0.019 for loom, p = 0.015 for 86% coherent, WRS; N = 3). Vertical dashed lines show the average time of jump. (**c**) Example extracellular recordings during jump experiments before and after ZD7288 appliCa_T_ion. The control responses displayed have 182 and 165 spikes, and responses after ZD7288 have 128 and 122 spikes for the standard loom and 86% coherent loom, respectively. (**d**) Rasters and instantaneous firing rates after injection of ZD7288 through the eye in restrained animals, reduced LGMD responses in a similar manner as intracellular or visually guided appliCa_T_ion (**Fig. 2i**), implying that the stereotactic injection method successfully targeted the LGMD.

## DISCUSSION

Here, we provide the first demonstration of selectivity to spatial coherence for an ecologically important escape behavior (**Fig. 1**). We also show that this spatial discrimination relies on processing within the dendrites of the LGMD. Whether the LGMD could discriminate between spatial patterns derived from a looming stimulus was previously unknown. Nor was it known how neurons could discriminate broad spatial patterns of inputs to implement object segmentation. To examine this issue, we provided the first characterization of active conductances within the dendrites of the LGMD and demonstrated that HCN and inactivating K^+^ channels produced selectivity for spatial coherence within the LGMD’s dendrites receiving precise retinotopic excitation (**Figs. 2 & 4**). Although our results demonstrate that spatial selectivity is in large part implemented within the LGMD’s dendritic arbor, they do not rule out possible additional presynaptic mechanisms supplementing those described here. Further, by blocking HCN channels in freely moving animals, we showed that the selectivity of escape behavior depends on the enhancement of spatially coherent responses due to HCN channels in the LGMD (**Fig. 6**).

Although our experimental data demonstrated that HCN channels produce a selective enhancement for inputs generated by spatially coherent approaching objects and that this leads to behavioral selectivity, there are no previously described mechanisms by which ion channels could produce this remarkable spatial selectivity. Based on extensive biophysical modeling, we developed a plausible hypothesis explaining the underlying mechanisms, schematically illustrated in **Fig. 7**. These mechanisms involve dendritic compartmentalization, regulation of membrane potential to control levels of K^+^ and Ca^2+^ channel inactivation, regulation of bursting, and competition between depolarizing and hyperpolarizing conductances within a compartmentalized dendritic arbor. While HCN channels play a critical role in these processes, each aspect depends on their interaction with other membrane conductances. We postulate that spatially incoherent visual stimuli generate spatially dispersed synaptic inputs which increase the activation of K_D-like_. This reduces the depolarization generated by synaptic inputs (**Fig. 4d,e; Supplementary Fig. 4**) and the K_D-like_ activation reduces its own slow inactivation. For spatially coherent stimuli, in contrast, the synaptic inputs combine with the tonic activity of g_H_ to maintain local depolarization sufficient to inactivate K_D-like_. The sustained depolarization propagates to the SIZ where it inactivates Ca_T_ channels, reducing burst spiking and negative feedback caused by K_Ca_. Conversely, incoherent stimuli generate more transient depolarization with less inactivation of K_D-like_ and Ca_T_ and more burst firing followed by K_Ca_ activation that further prevents sustained spiking. Escape jumps require a coordinated sequence of motor activation generated by a specific LGMD firing pattern and cannot be initiated by transient bursts of spiking^26,29^. While HCN channels often promote rhythmicity^37^, increases in bursting after removal of g_H_ due to de-inactivation of Ca_T_ channels has also been reported in cortical neurons and tied to absence epilepsy^47,48^.

**Figure 7.**
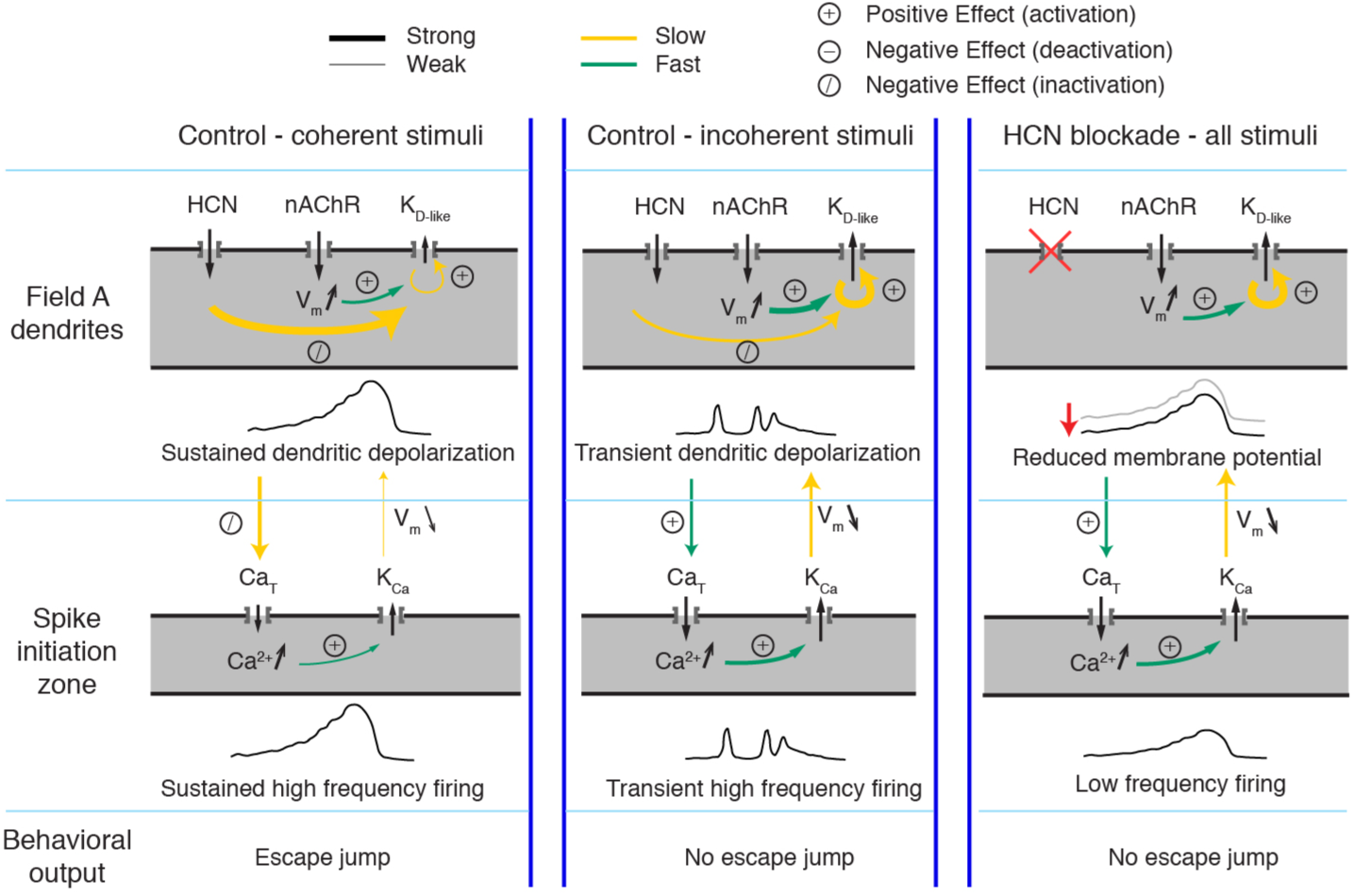
Schematic illustration of the effects of HCN channels on responses to looming stimuli. Arrows indicate current flow, changes in Ca^2+^ or membrane potential, and the strongest interaction between channels, dendrite and SIZ, during looming stimulus detection. For all visual stimuli, excitatory nicotinic acetylcholine receptors (nAChR) produce a fast activation of K_D-like_. In control conditions during coherent stimuli (left), the addition of g_H_ on top of nAChR activation produces sustained depolarization which slowly inactivates K_D-like_ channels within field A and Ca_T_ channels at the SIZ. This results in reduced K_D-like_ and K_Ca_ conductances and sustained high frequency firing. In control conditions during incoherent stimuli (middle), the increased dendritic area of nAChR activation increases the number of K_D-like_ channels activated. The resulting increase in the hyperpolarizing K_D-like_ conductance prevents the g_H_-dependent inactivation of K_D-like_ channels, producing only transient depolarization. The transient depolarization initiates Ca_T_ driven bursts and subsequent K_Ca_ conductance activation. This prevents the sustained high frequency firing necessary for initiating escape. After g_H_ blockade (right), the resting membrane potential is reduced, increasing the activatable K_D-like_ and Ca_T_ channels. Without g_H_, depolarization cannot be sustained high enough to inactivate these channels leading to an increased K_D-like_ and K_Ca_ conductances and lower frequency firing which fails to produce escape behaviors.

The model illustrated in Fig. 5 and 7 supports our experimental results and was the simplest that could be derived from them. Although the specific kinetics and distributions of several channels in the LGMD remain to be characterized, the model is well grounded in experimental data (**Material and Methods**). Further, extensive searches through the possible parameter space did not yield other combinations of mechanisms that reproduced the broad range of experimental data available to constrain the model. Yet, the model did not reproduce quantitatively all our experimental data: for example, it underestimated the amount of transient firing at high coherence and, vice-versa, overestimated its sustained firing (**Fig. 5e**; **Supplementary Fig. 5d,e**, dashed lines). One likely reason is that details of the bursting mechanisms may be imperfectly tuned in the model, due to the absence of a second calcium sensitive K^+^ conductance^35^ or still uncharacterized properties of an M current (unpublished observations). Further confirmation of this will require future experimental tests of channel properties predicted by the model, including the location of HCN and K_D-like_ channels within field A (**Fig. 2e**), that K_D-like_ and Ca_T_ inactivate above -70 mV, and that K_D-like_ inactivates slowly (in the range of 0.3-2 seconds).

This work also generates hypotheses about the role of the LGMD output in the production of escape behavior. Incoherent stimuli generated bursts of LGMD activity, but not the sustained firing of increasing frequency seen in response to coherent looming stimuli, and failed to initiate escape behavior (**Fig. 1f**). As the escape jump requires a coordinated sequence of muscle activation, this suggests that bursts interrupt the motor sequence before all the required phases are completed. One could think of a burst of spikes followed by a pause in firing as a type of ‘abort signal’ canceling the jump program before it reaches completion. Further behavioral tests, ideally with *in vivo* stimulation, could be used to test this hypothesis.

Additionally, the current work also suggests further reaching hypotheses. HCN channels have long been known to influence dendritic integration in hippocampal pyramidal neurons^40^, and KD has been tied to extended synaptic integration in the same neurons^45^. More recently, dendritic K^+^ channels have been found to compartmentalize the dendrites, and experiments and modeling have suggested that spatiotemporal interactions between HCN and K^+^ channels regulate neuronal excitability^49,50,41^. While the spatial pattern of synaptic inputs *in vivo* is unknown in pyramidal neurons, it is possible that a similar selectivity for spatial patterns arises by mechanisms analogous to those described here. In thalamocortical neurons, HCN channels influence K^+^ and Ca^2+^ channel inactivation, thereby regulating bursting and excitability^47,51^. HCN regulation of bursting has been tied to a rat model of absence epilepsy^48,52^ and may also contribute to human epilepsy^53^. In addition to possible disease states, HCN-dependent regulation of persistent or burst firing has also been involved in working memory^54^.

In summary, our results highlight how nonlinear dendritic conductances and the interactions between them can confer the ability to reliably select synaptic patterns appropriate for the generation of visually guided escape behaviors. This demonstrates the remarkable computing power of individual neurons and may help design object segmentation algorithms for bio-inspired collision avoidance systems. As HCN conductances are ubiquitous, they likely contribute to implement analogous computations in other species, including our own^53^.

## Acknowledgments

We would like to thanks Y. Zhu and E. Sung for the two-photon LGMD scan and reconstruction, respectively, and S. Peron and J. Reimer for feedback on an earlier draft of the manuscript. F.G. would like to thank the Marine Biological Laboratory (Woods Hole, MA) for a 2006 Faculty Summer Research Fellowship that helped initiate this line of research. This work was supported by grants from NIH (MH-065339) and NSF (DMS-1120952), as well as by a NEI Core Grant for Vision Research (EY-002520-37). This work was also supported in part by the Cyberinfrastructure for Computational Research, funded by NSF under Grant CNS-0821727 and Rice University.

## Methods

### Animals

All experiments were performed on adult grasshoppers 7-12 weeks of age (*Schistocerca americana*). Animals were reared in a crowded laboratory colony under 12 h light/dark conditions. For experiments preference was given to larger females ~3 weeks after final molt that were alert and responsive. Sample sizes were not predetermined before experiments. Animals were selected for health and size without randomization, and investigators were not blinded to experimental conditions.

### Surgery

The surgical procedure for intracellular recordings was described previously^30,32^. For extracellular DCMD recordings in freely moving animals, we developed a novel chronic implant technique, allowing the same animals to be recorded over many days, based on previous methods^24^. Grasshoppers were fixed ventral side up and a rectangular window was opened in their thorax. Air sacs were removed and the trachea were carefully separated to expose the ventral nerve cords. Two teflon-coated stainless steel wires 50 µm in diameter were cut to a length of ~4 cm and fashioned into hooks with the coating removed from the inside edge of the crook (supplier: California Fine Wire, Grover Beach, CA). The electrodes were implanted with the deinsulated region placed against the dorsomedial edge of the left nerve cord between the pro- and meso-thoracic ganglia. Slight tension was applied to the cord to maintain a fixed position against the wires, and the wires were set in place by waxing them to the left side of the thorax. The cuticle window was then closed and sealed with a wax-rosin mixture and Vetbond (3M, St. Paul, MN). A ground electrode made of the same wire as the hooks was placed outside on the thorax and embedded in the wax. All three wires were routed laterally and fixed to the dorsal pronotum using the wax-rosin mixture with just enough slack to allow normal pronotum movement. The ends of the wires were de-insulated and positioned pointing up to prevent the animal from reaching them. After the surgery, animals were allowed a day to recover, and survived for up to 7 months during which time the animals behaved normally.

To connect the electrode wires to the amplifier during an experiment, the animals were held in place with transparent surgical tape (Dukal Corp, Ronkonkoma, NY). The free ends of the implanted electrodes were each attached to polyurethane-coated hook-up wire with a pair of gold-plated miniature connectors (0508 and 3061, Mill-Max, Oyster Bay, NY; wire diameter: 160 µm or 34 AWG, Belden, St. Louis, MO). The hook-up wires were braided together and loosely suspended directly above the animal to allow unrestrained movement. Neither the implantation surgery or connection of implanted wires to the amplifier caused a significant reduction in escape behavior (**Supplementary Fig. 6b**).

### Visual stimuli

Visual stimuli presented during jump experiments were generated with custom software on a personal computer (PC) running the real-time operating system QNX 4 (QNX Software Systems), as previously described^24^. Identical visual stimuli for electrophysiological experiments were generated using Matlab and the PsychToolbox (PTB-3) on a PC running Windows XP. In both cases, a conventional cathode ray tube (CRT) monitor refreshed at 200 frames per second was used for stimulus display (LG Electronics). Both monitors were calibrated to ensure linear, 6-bit resolution control over luminance levels. Visual stimuli were presented in blocks with each stimulus shown once per block and the order within the block randomized by the stimulus software for all experiments. For wide field stimuli presented to restrained animals, a 90-120 s delay was used between stimuli and grasshoppers were repeatedly brushed and exposed to light flashes and high frequency sounds to decrease habituation. Some animals still exhibited pronounced visual habituation (> 50% reduction in peak firing rate from that animal's average response to the stimulus), and these data were excluded from analysis. In escape behavior experiments, a delay of at least 5 min (and usually ~15 min) was used between stimuli to prevent habituation.

Looming stimuli consisted of dark squares simulating the approach of a solid object on a collision course with the animal^23^. Briefly, the instantaneous angular size, *θ(t)*, subtended at one eye by a square of radius, *l*, approaching the animal at constant speed, *v*, is fully characterized by the ratio, *l*/|*v*|, since *θ(t)* = 2 tan^−1^ [*l*/(*vt*)]. By convention, *v* < 0 for approaching stimuli and *t* < 0 before collision. Stimuli simulated approach with *l*/|*v*| values of 50 or 80 ms from an initial subtended angle of 1.2° until filling the vertical axis of the screen (300 mm), lasting approximately 4 and 7 s for *l/|v|* = 50 and 80 ms, respectively. The maximum *θ* values reached by the stimuli were either 136° or 80° for the freely behaving or restrained preparations, respectively, due to the differing distances of the eye to the screen.

‘Coarse’ looming stimuli were generated as in our earlier work^32^. Briefly, the stimulation monitor was first pixelated with a spatial resolution approximating that of the locust eye (2-3° x 2-3°), referred to as ‘coarse’ pixels. Each coarse pixel’s luminance followed the same time course as that elicited by the edge of the simulated approaching object sweeping over its area. To alter the spatial coherence of these stimuli, a random two-dimensional Gaussian jitter with zero mean was added to each coarse pixel screen location. The jittered positions were rounded to the nearest available coarse pixel location on the screen to prevent any coarse pixels from overlapping. The standard deviation of the Gaussian was altered between 0 and 80° to control the amount of shifting and thus the resulting spatial coherence of the randomized stimulus.

For localized light flashes, a 1° × 1° luminance increase was presented briefly (~1 s) on a black background in the dark (**Supplementary Fig. 1a-c**). A window of 200 ms following the flash onset was used to quantify LGMD activity.

### Escape behavior

The behavioral experiments were conducted as previously described^35^. They were recorded with a high-speed digital video camera (GZL-CL-22C5M; Point Grey, Richmond, BC, Canada), equipped with a variable zoom lens (M6Z 1212-3S; Computar, Cary, NC). Image frames were recorded at 200 frames per second with the acquisition of each frame synchronized to the vertical refresh of the visual stimulation display (Xtium-CL PX4; Teledyne Dalsa, Waterloo Canada). Videos were made from the images and saved in lossless motion JPEG format using custom Matlab code. Measurements of the stimulus coherence's effect on escape behavior (**Fig. 1f**) include a total of 202 trials from 66 animals with 1-9 trials per animal. Animals which did not jump in response to any stimuli were excluded from analysis, as done previously^35^.

### Electrophysiology

Electrophysiological experiments were performed as described previously ^30,32^. Briefly, sharp-electrode LGMD intracellular recordings were carried out in both voltage-clamp and current-clamp modes using thin walled borosilicate glass pipettes (outer/inner diameter: 1.2/0.9 mm; WPI, Sarasota, FL). After amplification, intracellular signals were low-pass filtered (cutoff frequency: 10 kHz for V_m_, and 5 kHz for I_m_) and digitized at a sampling rate of at least 20 kHz.

We used a single electrode clamp amplifier capable of operating in discontinuous mode at high switching frequencies (typically ~25 kHz; SEC-10, NPI, Tamm, Germany). Responses to visual stimulation were measured in bridge mode, current injections were applied in discontinuous current clamp mode (DCC), and voltage-clamp recordings in discontinuous single-electrode voltage-clamp mode (dSEVC). All dSEVC electrodes had resistances <15 MΩ. Electrode resistance and capacitance were fully compensated in the bath immediately prior to tissue penetration and capacitance compensation was readjusted after entering the neuron. If capacitance could not be fully compensated the recording was not used. In addition to previously described methods, a fluorescent dye (Alexa Fluor 594 hydrazide salt; Molecular Probes) was injected intracellularly and the cell was imaged with a CCD camera mounted to a stereomicroscope (GuppyPro F125B; Allied Vision Technologies, Exton, PA). This allowed subsequent visually guided positioning of the recording electrode.

During voltage clamp recordings, the membrane potential and current were measured simultaneously to ensure the desired membrane potential was maintained at the electrode location. The LGMD neuron is not electrotonically compact^55^ and therefore the issue arises of how well its dendritic membrane potential is controlled through voltage-clamping at a single location (“space clamp”). The quality of the space clamp cannot be measured with a single electrode recording. In pyramidal neurons, the steady-state dendritic membrane potential is largely uncontrolled when voltage clamping originates at the soma^56^. In contrast, in Purkinje cells, which have a dendritic structure more closely resembling that of the LGMD, the steady-state dendritic membrane potential is well controlled from the soma^57^. Simulations in NEURON (details below) were used to estimate the quality of the space clamp in the LGMD. For electrodes placed at the base of field A the average steady-state change in membrane potential within field A was 95% of the desired change (i.e., starting at rest, −65 mV, a −30 mV step to −95 mV, yielded an average membrane potential across field A of −65 + 0.95 · (−30) or −93.5 mV). For electrodes placed further away from the base of field A, the quality of the space clamp decreased. Across the dendritic region used for voltage clamp recordings, the estimated quality of the space clamp ranged from 83-95% (average voltage command attenuation of 5-17%).

For characterizing g_H_, 1-2 s hyperpolarizing current or voltage steps were injected in DCC or dSEVC mode, respectively, with 5 s between steps. Different step amplitudes were randomly interleaved and at least 6 trials per step amplitude per animal were acquired. For each recording, we used at least 4 step amplitudes, with values selected to cover the activation range of g_H_. Example recordings are shown in **Supplementary Fig. 1a**. In most experiments, no holding current was applied between steps (held at the resting membrane potential), while in some experiments a positive holding current was applied (held near -50 mV) before the hyperpolarizing step and in other experiments a negative holding current was used (held near −115 mV) with depolarizing steps. Estimated activation curves (see below) were not different for recordings with different holding potentials, and the data were combined for analysis. Extracellular recordings were taken between the two hook electrodes on the nerve cord, differentially amplified and bandpass filtered from 100 to 5000 Hz (A-M Systems, model 1700, Carlsborg, WA). The amplitude of DCMD spikes was consistently the largest, allowing their identification with a simple threshold. DCMD spikes uniquely identify the LGMD neuron as they are in one-to-one correspondence with those of the LGMD^19^.

Experiments with hyperpolarizing current during visual stimulation (**Supplementary Fig. 3b**) were conducted by first staining the LGMD and then inserting an electrode near the base of field A. Visual stimuli of different coherences were presented either with zero current or −2.5 nA current injected from 20 s before stimulus onset through the end of the stimulus. Sets of single trials of each stimulus (randomized) were presented while alternating between 0 and −2.5 nA currents and continued until at least 3 trials of each stimulus were presented for both conditions.

## Pharmacology

Drugs were prepared in aqueous solution and mixed with physiological saline containing fast green (0.5%) to visually monitor the affected region. They were puffed using a pneumatic picopump (WPI, PV830). For restrained experiments, injection pipettes had tip diameters of ~2 µm and were visually positioned with a micromanipulator against the posterior edge of the lobula, close enough that the ejected solution penetrated the optic lobe. Drugs were gradually applied while monitoring responses of the LGMD to visual inputs, and care was taken to prevent spread into presynaptic neuropils. Additionally, saline in the bath was exchanged immediately after puffing to prevent diffusion to other brain areas. We used drug concentrations of 10 mM for both ZD7288 and 4AP in the extracellular puff pipette. These concentrations were adjusted in pilot experiments to account for the low mobility of the drugs through the tissue *in vivo*, taking into account their approximate final concentration, as explained below.

Due to dilution of the drugs in the saline bath after puffing, the exact drug concentration at the level of the LGMD cannot be determined. However, our best estimate is ~200 µM for both ZD7288 and 4AP. This estimate comes from comparing the effect of the puffed drugs to those observed after bath application of the same drugs. For example, when bath applying ZD7288, the same level of blockade as from local puffing was achieved by adding 100 µl of 20 mM drug to ~5.5 ml of bath saline for a final concentration of ~350 µM. This concentration is an upper bound on the concentration at the level of the LGMD, since it lies ~150 µm deep within the optic lobe. For local puffing, less than 1 µl of drug was used, which would generate a final bath concentration well below 1 µM after exchanging the saline in the bath, as explained above.

For intracellular application, the drug concentrations in the pipette were 1-5 mM for ZD7288 and 5 mM for 4AP. The final concentration inside the LGMD cannot be determined but is likely considerably lower, due to the large volume of the cell and the submicron diameter of the pipette. In those experiments, the effects of the drugs were comparable to those observed with extracellular application.

Although it cannot be known whether intracellularly applied ZD7288 or 4AP diffused out from within the LGMD, this seems highly unlikely to have affected our results. For example, the effects of intracellular ZD7288 application on the LGMD's membrane properties were consistent from a minute to an hour after application, giving no evidence of a slow diffusion across tissue that may have affected presynaptic sites. Further, the membrane effects on the LGMD were the same whether excitatory synaptic inputs were blocked with mecamylamine or not. To further rule out the possibility that the effects of ZD7288 observed during visual stimulation were caused by diffusion to presynaptic sites after intracellular application, we conducted visual stimulation experiments in which the g_H_ conductance was blocked intracellularly with Cs^+^ (**Supplementary Fig. 1 & 3**). Similar effects were seen on visual responses compared to ZD7288 application, although Cs^+^ was not as specific a blocker since there was also evidence of partial block of K^+^ conductances. For these experiments, a concentration of 150 mM CsCl was used in the recording pipette. We also attempted to block intracellularly the inactivating K^+^ conductance by using 4-aminopyridine methiodide (4APMI) which is membrane impermeant^58^. 4APMI reacted strongly with the silver wire in the electrode forming AgI crystals, so a platinum wire was used for the experiments. Unfortunately, 4APMI which is larger than 4AP failed to block the inactivating K^+^ conductance even at recording pipette concentrations as high as 50 mM. Nonetheless, presynaptic effects are unlikely as we never observed increases in spontaneous EPSPs within the LGMD following 4AP application and the presynaptic neurons have no information about the overall spatial pattern of the stimulus. In all there were no indications of any nonspecific drug effects on presynaptic neurons that might have influenced visual responses.

To observe the effects of ZD7288 in freely moving animals, stereotaxic injections were made through a hole in the dorsal rim region of the right eye. The animal was restrained and the head was placed in a small clamp attached to a 3-axis micromanipulator (Narishige, Japan). Head tilt was positioned manually by fixing the animal at the pronotum. After the head was precisely positioned, a ~0.5 mm hole was made through the dorsal end of the eye with a steel probe. A drop of saline solution was placed covering the hole to prevent drying or coagulation of the hemolymph. A glass pipette with a tip diameter of 1-2 µm and a taper length >2 mm from shoulder to tip was positioned with a Leica manual micromanipulator and lowered just above the eye. The ZD7288 solution (2 mM in saline with 1% fast green) was puffed into the saline drop covering the dorsal rim to determine the appropriate air pressure ejection level. The saline droplet was immediately removed and replaced to prevent spread of ZD7288 to photoreceptors. Next, the pipette was lowered through the eye along the dorsal rim of the optic lobe to the lobula (~1.5 mm) while enough positive pressure was maintained to prevent clogging. In control experiments, LGMD activity was measured before and after penetration of the pipette in the lobula to ensure that visual inputs were not damaged by the procedure (**Supplementary Fig. 6c**). Ejection volume was estimated from monitoring changes in the meniscus position of the saline within the visible region of the pipette. After pressure ejection of ZD7288, the pipette was removed and checked for clogs or breaks. The hole in the eye was sealed with a small amount of Vetbond (3M), carefully ensuring that no glue spread onto the rest of the eye.

Following the conclusion of the experiment, the animal was euthanized and the head was dissected (~2 hours post injection). Fast green staining was used to confirm that the solution was injected into the lobula (**Supplementary Fig. 6d**). In initial experiments, bath application of ZD7288 was found to reduce visual responses as did application of ZD7288 directly to photoreceptors. When puffing ZD7288 within the lobula, however, even if the solution occasionally spread to the medulla or lamina visual responses remained similar to those observed after intracellular application. This suggests that there are likely HCN channels within the photoreceptor layer, as is the case in mammals^59^, but that any HCN channels within the medulla and lamina^60^ do not influence LGMD inputs under our experimental conditions. Because ZD7288 was applied extracellularly, it may have affected other descending neurons whose processes are located in the immediate vicinity of the LGMD dendrites. This is however unlikely to have affected escape behaviors, since decrease in escape was tightly correlated with a reduction of LGMD firing rate determined in independent experiments (**Fig. 2l**). In addition, earlier selective ablation experiments have shown that under our experimental conditions nearly all escape behaviors depend solely on LGMD firing^28^.

## Data analysis and statistics

Data analysis was carried out with custom MATLAB code (MathWorks). Linear fits were based on Pearson's linear correlation coefficient, denoted by 'r' in figure legends. Non-linear fits, including the activation curve and time constant in Figure 2 and all exponential fits described below were made with the Matlab function ‘lsqcurvefit’, which minimizes the least square error between the data and fitting function. Goodness of fit was denoted by R^2^, calculated as one minus the sum squared error of the fit divided by the sum square deviation from the mean of the data.

The sag amplitude was measured as the difference in membrane potential between the peak hyperpolarization during a current step and the steady-state value at the end of the step. The sag time constant was calculated from fitting a single exponential to the membrane potential for the period starting 15 ms after peak hyperpolarization to the end of the current step (**Supplementary Fig. 1a**). The hyperpolarizing step currents were also used for calculating membrane time constants. The membrane time constant was calculated by fitting a single exponential to the membrane potential for the period from 0.5 to 13 ms after the start of hyperpolarizing current injection.

The fitted activation curve of the HCN conductance was based on a Boltzmann equation reflected along the voltage axis to produce decreasing *g*_*H*_ with increasing *v*:

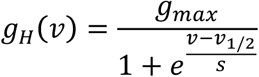

The steady-state conductance, *g*_*H*_ is a function of the membrane potential, *v*, depending on three parameters: the maximum conductance, *g*_*max*_ the half-activation potential, *v*_1/2_ and the steepness, *s*. The parameters were fitted from voltage-clamp data based on the equation

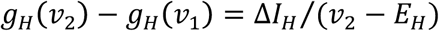

where *v*_*1*_ and *v*_*2*_ are the starting and ending clamp potentials and *E*_*H*_ is the reversal potential of the HCN conductance, -35 mV, used for all animals. Δ*I*_*H*_ is the experimentally measured change in membrane current produced by the voltage step after transients have settled. Δ*I*_*H*_ was measured by fitting a single exponential to the current time-course 15 ms after the step onset and up to its end (**Supplementary Fig. 1a**). This period captured the slow change in clamp current due to *gH* and offered clear experimental advantages over other estimations methods. As all experiments were done *in vivo*, it was not feasible to reliably block other putative voltage-gated channels. Hence, the most reliable measurement of Δ*I*_*H*_ were obtained at hyperpolarized membrane potentials where other active conductances can be safely discounted. Voltage clamping the LGMD to depolarized potentials where all HCN channels will be closed (> -40 mV) was not technically feasible, and the use of tail currents yielded less reliable measurements due to contamination by other active conductances.

The time constant of the HCN conductance (τ_H_) was fit using a function symmetric with respect to its maximum, τ_*max*_,

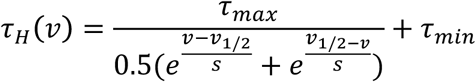

Here, *v*_*1/2*_ is the membrane potential with the slowest activation, *s* is the steepness, and τ_*min*_ the minimum activation time. Fitted points were obtained from the single exponential fits to I_H_ for both hyperpolarizing (channel opening) and depolarizing (channel closing) voltage steps.

Comparisons of sag amplitudes were obtained with current steps yielding a peak hyperpolarization of ~105 mV (**Fig. 2c-e**). For **Figure 2c,d**, all values to steps within the range of -95 to -115 mV were pooled. For **Figure 2e**, interpolation of values at nearby potentials was used to estimate sag amplitude at -105 mV to have a single common value for all recordings. Statistical comparisons between sag measurements in different subcompartments of the LGMD (**Fig. 2c,d**) were carried out using a Kruskall-Wallis analysis of variance (ANOVA) corrected for multiple comparisons with Tukey's Honestly Significant Difference Procedure (KW-MC). To determine the correct statistical test for comparison we used a Lillifors test of normality (alpha = 0.20) and comparison of equality of variance. Much of the data was non-normally distributed and variances increased after drug application so most comparisons were made using the Wilcoxon rank sum test (WRS) which does not assume normality or equality of variance. For displaying non-normal data, average values were given as median and variance was displayed as median average deviation (mad). Mean and standard deviation were used for normally distributed values, as indicated in figure legends. Percent activation at rest (**Fig. 2h**) was calculated through bootstrapped activation curves from current clamp data. Unpaired t statistics were calculated from the bootstrapped mean and variance of activation at the resting membrane potential (-65 mV)^61^.

Simulated excitatory postsynaptic potentials (sEPSPs) were generated by injecting a series of five current waveforms with a set delay between them. Each waveform, *I(t)*, had a time course resembling that of an excitatory synaptic current,

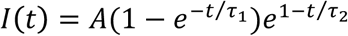

with peak amplitude *A*, rising time constant *τ*_*1*_ = 0.3 ms, and falling time constant *τ*_*2*_ = 3.0 ms. Summation was calculated as the ratio (p_5_-p_1_)/p_1_, with p_1_ and p_5_ being the peak amplitude of the membrane potential relative to rest during the 1^st^ and 5^th^ sEPSP. In **Fig. 3f** we plotted the integrated membrane potential (relative to rest) divided by the integrated input current (charge) giving a value in units of mV ms / nA ms = MΩ that is readily comparable to input resistance.

Normalized firing rates (**Fig. 2j** and **Fig. 4b**) were calculated by dividing the response amplitude for each stimulus by that animal's maximal response amplitude under control conditions (insets of **Fig. 2j** and **4b**). These individually normalized rates were then averaged across animals. The relative change in response due to a drug (**Fig. 4c**) was calculated by dividing the absolute difference in response between control and drug conditions by the lesser of the two firing rates (control in the case of 4AP, and post drug in the case of ZD7288). This produced percentages covering similar ranges, and so allowed for the best comparison and graphical illustration of their relative effects.

‘Sustained firing’ was defined as the longest period in which the instantaneous firing frequency remained above a 20 Hz threshold. For each trial, the number of spikes within this longest period was considered the ‘sustained response’ and all spikes outside of this period were counted as the ‘transient response’ (**Supplementary Fig. 5e,f**). These 'sustained' and 'transient' measures were used instead of 'burst' and 'non-burst' statistics based on interspike intervals because the LGMD can generate sustained high frequency firing with similar interspike intervals within and outside of bursts.

To compare changes in membrane potential and stimulus angular distance (**Fig. 4d; Supplementary Figure 4**), we identified newly changing coarse pixels in a specified stimulus frame from those that had begun to darken from their background luminance value in earlier frames (‘earlier changing’). We then computed the mean minimal distance of newly changing coarse pixels with respect to earlier changing ones. In parallel, changes in the membrane potential were averaged from 25 ms following the appearance of the newly changing coarse pixels until a new group of pixels began to darken. More precisely, we identified six time periods during the stimuli when the luminance of newly changing coarse pixels is decreasing for over 50 ms and they typically have mean angular distances larger than 1° from earlier changing ones. For these six different time periods during each trial, we calculated the linear correlation between these mean angular distances and membrane depolarizations, as explained above. Early in the stimulus presentation, there are fewer coarse pixels changing luminance and less resulting depolarization. To better illustrate the relationship between these variables, the angular distances and membrane potentials were normalized independently for each of the six time windows. The normalization consisted of subtracting the minimum control value and then dividing by the range for control data within each time window. The unnormalized data and example stimulus frames from all time periods are shown in **Supplementary Figure 4**.

To compare jump probabilities between saline- and ZD7288-injected animals (Fig. 6a), we computed 95% bootstrap confidence intervals of the population mean in each condition with the help of the built-in Matlab function ‘bootci’ (using the bias corrected and accelerated method). If there was no overlap of the 95% confidence intervals, the groups were considered significantly different. The reported p-values for these comparisons were the ‘achieved significance level’ (ASL) statistic for two-sample testing of equality of means with unequal variance (Algorithm 16.2 in ref. *61*).

For calculation of the percent spatial coherence of a stimulus, we determined the minimal total angular distance that coarse pixels of a spatially altered stimulus had to be moved to produce an unaltered coarse looming stimulus. This distance was then normalized by the angular distance between a coarse loom and one with completely random spatial positions. This computation yields coherence values ranging from 0% for random spatial positions to 100% for coarse and standard looming stimuli. Coherence selectivity was calculated as the slope of the relationship between stimulus coherence and spike count and is reported in units of spikes per percent coherence. For control experiments the median correlation coefficient of this relationship between stimulus coherence and spike count was 0.97, making the regression slope a reliable indicator of the selectivity.

### Neuronal modeling

To better understand the mechanisms of the LGMD's remarkable coherence selectivity, we developed a detailed biophysical model using the NEURON simulation environment. We employed the parallel version of NEURON and a Rice University supercomputing cluster for extensive parameter sweeps and simulations. Three dimensional reconstructions of the LGMD's dendritic morphology were obtained from 2-photon scans using the software suite Vaa3D (vaa3d.org). The resulting model contained 2518 compartments, 1266 of which belonged to dendritic field A.

To reproduce the active properties of the LGMD several voltage-gated channel types were included. Some of them had been used in previous simulations^55,62^, including the fast Na^+^ and delayed rectifier K^+^ (KDR) channels generating action potentials. KDR channels were distributed throughout the cell, but dendritic branches contained no fast Na^+^ channels as supralinear summation is never seen in LGMD dendrites. HCN channels had kinetics matching experimental data (**Fig. 2**) and were placed in dendritic field A with density increasing towards the distal dendritic endings. Inactivating K^+^ channels (K_D-like_) were also distributed throughout field A with density increasing toward distal endings. A slow non-inactivating K^+^ channel (M) was distributed throughout the axon, the spike initiation zone (SIZ), and the main neurite connecting the dendritic subfields to the SIZ. Its peak density was at the SIZ. Additionally, low-threshold Ca^2+^ (Ca_T_) and Ca^2+^-dependent K^+^ (K_Ca_) channels were placed at the SIZ and on half of the neurite connecting the SIZ and the dendritic subfields, matching results from our earlier work^43^.

Effective modeling often relies on keeping things as simple as possible, so we initially tested a previous LGMD model^32^ with additional HCN channels matching experimental kinetics (**Fig. 2**) added to the dendrites. When this failed to reproduce any coherence selectivity, K_D-like_ channels were added. Then we added a complex dendritic morphology, complex presynaptic transforms, and additional active conductances to the model. While a wide range of parameters of this more complex model reproduced responses to current injection data, only one narrow parameter regime was found that reproduced the roles of g_H_ and K_D-like_ in the spatial coherence preference. The resulting model and mechanistic explanation (**Fig. 5** & **7**), while quite complex, is still the simplest model that reproduced the wide range of LGMD responses tested. Evaluation of how well this model informs about the actual neural processes requires some review of the experimental data to which it was constrained.

The strength and timing of synaptic inputs was generated based on single facet stimulation data^32^. Excitatory synaptic input locations were based on the retinotopy and synaptic overlap determined by functional imaging^30,31^. The time course of synaptic inputs was based on experiments stimulating individual facets^32^ and the pattern of depolarization measured during the current experiments. The presence of standard Hodgkin-Huxley Na^+^ and K^+^ currents was assumed, HCN and K_D-like_ channels were based on the current work, the Ca_T_ and K_Ca_ channels were based on ref. 43, the M current was based on our own currently unpublished findings. For each of these channels, conductance and kinetic parameters were adjusted to match experimental data with firing frequency vs. injected current curves and spike waveform used to tune fast Na^+^ and K^+^ channels, changes in input resistance and resting membrane potential after pharmacological blockade used to adjust HCN, K_D-like_, and M parameters, while Ca_T_ and K_Ca_ were adjusted to match intrinsic burst (currently unpublished) and spike frequency adaptation data^43^. Channel distributions were similarly grounded in experimental data when available, and were manually fit to find working parameters.

To estimate space clamp quality (see Methods: Electrophysiology), we used the Impedance object class in NEURON and measured the percent voltage attenuation from an electrode location to each compartment within field A of the model. The average attenuation was calculated by weighting each section by its surface area to calculate the average change of membrane potential within field A.

### Data and Code Availability

The full model and simulation code are available in the public repository ModelDB, accession number 195666. The experimental data and analysis code generated during the current study are available from the corresponding author on request.

